# Genomic Retargeting of Tumor Suppressors p53 and CTCF Promotes Oncogenesis

**DOI:** 10.1101/703124

**Authors:** Michal Schwartz, Avital Sarusi Portugez, Bracha Zukerman Attia, Miriam Tannenbaum, Olga Loza, Aliza Chase, Yousef Turman, Tommy Kaplan, Zaidoun Salah, Ofir Hakim

**Affiliations:** The Mina and Everard Goodman Faculty of Life Sciences, Bar-Ilan University, Ramat-Gan, Israel; School of Computer Science and Engineering, The Hebrew University of Jerusalem, Jerusalem, Israel; Al-Quds-Bard College for Arts and Sciences, Al-Quds University, Abu Dis, Palestinian Terretories; Department of Molecular Genetics, Weizmann Institute of Science, Rehovot, Israel

## Abstract

Gene transcription is substantially regulated by distant regulatory elements via combinatorial binding of transcription factors. It is more and more recognized that alterations in chromatin state and transcription factor binding in these distant regulatory elements may have key roles in cancer development. Here we focused on the first stages of oncogene induced carcinogenic transformation, and characterized the regulatory network underlying transcriptional reprogramming associated with this process. Using Hi-C data, we couple between differentially expressed genes and their differentially active regulatory elements and reveal two candidate transcription factors, p53 and CTCF, as major determinants of transcriptional reprogramming at early stages of HRas-induced transformation. Strikingly, the malignant transcriptional reprograming is promoted by redistribution of chromatin binding of these factors without major variation in their expression level. Our results demonstrate that alterations in the regulatory landscape have a major role in driving oncogene-induced transcriptional reprogramming.

## Introduction

Altered gene expression programs are a major factor in the development and expansion of cancer cells. These aberrant transcription patterns support and promote biological processes that are required along tumorigenesis, such as proliferation, resistance to cell death and induction of angiogenesis ^1^. Transcriptional programs are driven by a complex network of transcription factors acting on regulatory DNA elements such as enhancers (reviewed in^2^). Although changes in transcriptional programs during tumorigenesis could be driven directly by DNA mutations, in many cases they are affected indirectly by various altered cellular pathways. Transcription factors may affect transcriptional programs due to changes in their DNA binding, without harboring mutations, or change in their expression levels. Much of the effort to characterize the pathways contributing to cancer development mostly focus on identification of either upstream mutations ^3^ or downstream RNA and protein expression patterns of cancer cells ^4^. These are indeed very powerful approaches to reveal important drivers of cancer development and characterize cancer types. However, demarcating the intermediate molecular events triggering gene expression programs holds great promise for generating novel and improved diagnostic tools and identifying relevant targets for therapeutic interventions ^5,6^. Indeed, a recent pan-cancer analysis demonstrated global enhancer activation in most cancer types compared with matched normal tissues ^7^. Thus studying the regulatory basis of transcription reprogramming associated with carcinogenesis can define a set of transcription factors that are involved in shaping the transcriptional landscape of the cancer cell.

Transcription programs are regulated by a complex networks of transcription factors, cofactors and chromatin regulators ^2,8^. This regulation occurs via binding of transcription factors to regulatory elements, i.e. enhancers, in a sequence specific manner and recruitment of transcriptional coactivators or corepressors ^2,8^. Enhancer sequences contain multiple binding sites for a variety of transcription factors, and mediate transcriptional control by their combinatorial binding ^2,9^. This regulation is further modulated by the epigenetic status of enhancers. Therefore, mapping active enhancers can be key to delineating regulatory networks driving transcriptional reprogramming during the process of carcinogenesis. Active enhancer regions are associated with different molecular characteristics that allow their identification, such as histone modifications, specifically, H3K27ac and H3K4me1 ^10^ and occupancy of transcriptional co-activators, such as Ep300 ^11^. Importantly, for an enhancer to be bound by transcription factors and other regulatory DNA binding proteins, it must be accessible, therefore, a technique that is commonly used to map active regulatory elements is detection of open chromatin by sensitivity to nucleases ^12,13^. ATAC-seq is a recently developed sensitive method to measure chromatin accessibility which enables mapping of active regulatory regions in a genome-wide manner ^14^.

A significant challenge however, for inferring mechanistic principles of transcriptional regulation, is the difficulty to assign specific regulatory elements to their distant target genes. Enhancers and their regulated target genes can be located hundreds of kbs apart ^15^. Importantly, the spatial organization of genomic information is non-random and is a major regulatory component of gene transcription ^16,17^. In recent years it was demonstrated that mammalian chromosomes are partitioned into units of internal high spatial connectivity ^18^. These units, termed Topologically Associating Domains (TADs), are large (few tens of kb to 3 Mb) chromosomal units encompassing multiple genes and regulatory elements ^19^. Although the borders between TADs are considered cell-type invariant, at the local scale, chromosomal contacts between genes and their regulatory elements (within TADs) show cell-type specificity ^16^. Moreover, variation in gene expression within TADs is highly correlated during differentiation and response to external stimuli, suggesting that TADs insulate the activity of regulatory elements to specific genes ^15,20,21^. Therefore, TADs offer a proper and applicable framework to couple between regulatory elements and their target genes and study dynamic transcriptional regulation.

Here we combined, transcriptome analysis, genome-wide mapping of active regulatory sites with chromosome topology profiles to delineate the regulatory network underlying transcriptional reprogramming during carcinogenic transformation induced by over expression of oncogenic HRas in human mammary epithelial cells. Our results reveal major gene expression changes accompanied by significant alterations in the regulatory landscape upon this transformation process. Linking, in 3D, differentially expressed genes to the catalog of dynamic regulatory elements revealed two candidate transcription factors, p53 and CTCF, as major determinants of transcriptional reprogramming at early stages of HRas-induced transformation. Strikingly, the malignant transcriptional reprograming is promoted by redistribution of chromatin binding of these factors without major variation in their expression level. Moreover, this work highlights the power of the approach by which the transcriptional regulatory landscape can be analyzed in view of the 3D organization of the genome to reveal dynamic regulatory networks underlying cancer phenotypes.

## Results

Mutations in the HRas oncogene are among the most frequent in human tumors. To investigate transcriptional regulatory changes during early carcinogenesis, we introduced an oncogenic copy of HRas carrying a single point mutation at amino acid 12 (G12V) into MCF10A cells, a nontransformed, near-diploid, immortalized mammary epithelial cell line. These cells (from here on termed G12V MCF10A) first undergo growth arrest (data not shown) but after a few weeks continue growing and exhibit higher proliferation rates and higher cell survival capacity compared to their parental MCF10A counterparts (Fig. 1A,B). The tumorigenicity of the G12V MCF10A cells was tested by different means. Anchorage independent survival and growth are hallmarks of carcinogenic cells. Indeed, G12V MCF10A cells exhibit growth in soft agar (Fig. 1C) and adherence independent survival (Fig. 1D). Following HRas dependent transformation we noticed a morphological change towards a more mesenchymal morphology (data not shown), we therefore tested invasive potential and found that the G12V MCF10A cells harbor invasive properties as tested by Matrigel invasion assay (Fig. 1E). Finally, examination of their tumorigenic potential *in vivo* show that G12V MCF10A cells are able, although with low penetrance, to form small tumors when injected into mammary fat pad of Nod-SCID mice (Fig. 1F). Thus the introduction of G12V HRas oncogene into MCF10A rendered these cell tumorigenic.

**Fig.1.**
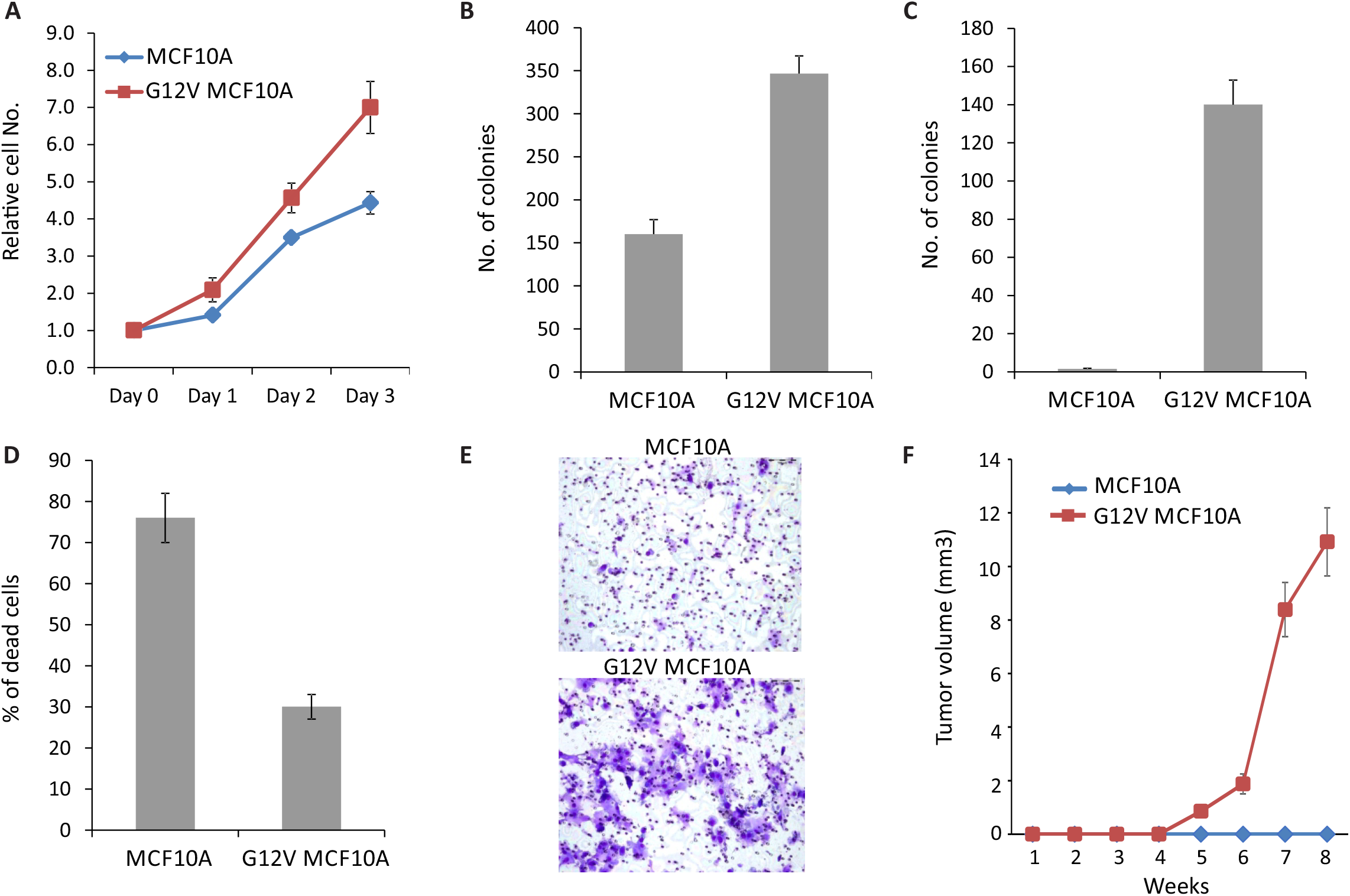
Characterization of G12V MCF10A cells. **(A).** Cell proliferation was measured by XTT assay. The relative number of cells, compared to day 0 is presented for MCF10A (blue line) and G12V MCF10A cells (red line). Error bars represent standard deviations of three independent replicates. **(B).** Cell survival was measured by colony formation assay. The number of colonies counted after two weeks of MCF10A and G12V MCF10A cells are presented. Error bars represent standard deviations of three independent replicates. **(C).** Anchorage independent growth was measured by soft agar assay. The number of colonies formed for MCF10A and G12V MCF10A cells is presented. Error bars represent standard deviations of three independent replicates. **(D)**. Resistance to anoikis was measured by nchorage independent cell death assay. The percentage of cell death for MCF10A and G12V MCF10A cells is presented. Error bars represent standard deviations of three independent replicates. **(E).** Representative images of Boyden Chamber Matrigel invasion assay of MCF10A and G12V MCF10A cells**. (F).** Tumor growth curve of mammary fat pad tumors in Nod-SCID mice injected with either MCF10A or G12V MCF10A cells (n=6mice/group). Error bars represent standard deviations.

In order to define the transcriptional reprogramming induced by the introduction of G12V HRas oncogene in MCF10A cells, we performed RNA-sequencing comparing G12V and parental MCF10A cells. We identified 1144 up-regulated and 1136 down-regulated genes (>1.3 fold, p<0.05) accompanying this transformation process (Fig. 2A and Supplementary table 1). Down-regulated genes are enriched in functions of cell death (p=2.67E-18) and apoptosis (p=2.53E-16) (Fig. 2B). Up-regulated genes are enriched in functions of invasion (p=2.5E-50), cell survival (p=1.42E-26), and cell movement (1.64E-25) and include tumor related genes (Fig. 2C). The enriched molecular pathways (Fig. 2D) also include some signaling pathways tightly related to breast cancer, such as IL-8 signaling ^22^ and p53 signaling ^3,4^. The top enriched upstream regulatory factors (Fig. 2E) also include p53. Thus the transcriptional reprogramming accompanying the expression of the G12V HRas oncogene is indicative of tumorigenic process.

**Fig. 2.**
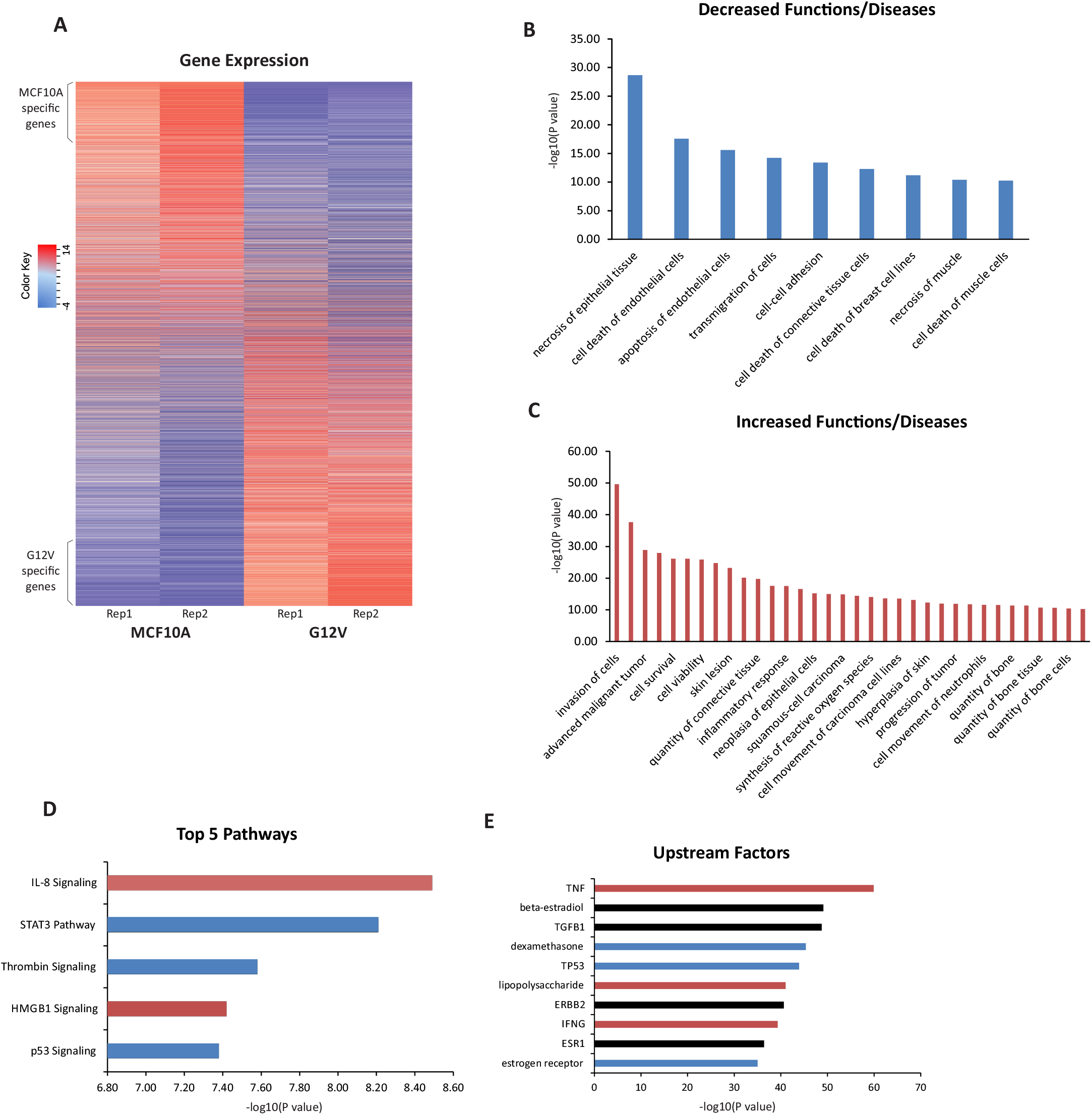
Transcriptional reprograming induced by the G12V HRas oncogene. **(A)** Heatmap showing gene expression of genes in MCF10A and G12V MCF10A cells from two replicas. Rows were first ordered based on log2 fold change and then by expression value. **(B-C)** Functions and Diseases enriched in down-regulated genes (B) and up-regulated genes (C) with a log10 pVal >10. **(D)** Top five significantly affected pathways according to the DE genes. **(E)** Top ten upstream regulators which can explain the changes in gene expression. Red - Predicted activation; Blue-Predicted inhibition; Black-No specific direction of activity. Analysis in B-E was done using the Ingenuity Pathway Analysis software.

In order to examine the changes in regulatory chromatin that dictate transcriptional reprogramming we applied ATAC-seq which allows discovery of accessible chromatin loci genome-wide. We identified 42,546 accessible loci in MCF10A parental cells and 46,367 in G12V MCF10A cells. Using stringent analysis of differential peaks in MACS we identified a few thousands of increased and decreased peaks representing Differentially Accessible Regions (DARs, Fig. 3A, B). 5,355 were induced in G12V MCF10A compared to parental cells (gained DARs) while 7,589 were reduced (lost DARs). Overall, the distribution of accessible regions in the genome (Fig. 3C) is similar to what was previously reported ^13^ with ~30% of loci near promoters. Interestingly the proportion of DARs at promoters is much lower (5% in G12V and 8% in MCF10A) relative to general distribution, indicating that the major changes in regulatory elements are at gene-distant regions (Fig. 3C). Examination of the connection between chromatin accessibility at promoters and gene expression reveals that up-regulated genes are more associated with gained accessibility while down-regulated genes associate more with loss of accessible regions (Fig. 3D). This reflects directional association between changes in gene expression and changes in their promoter accessibility.

**Fig. 3.**
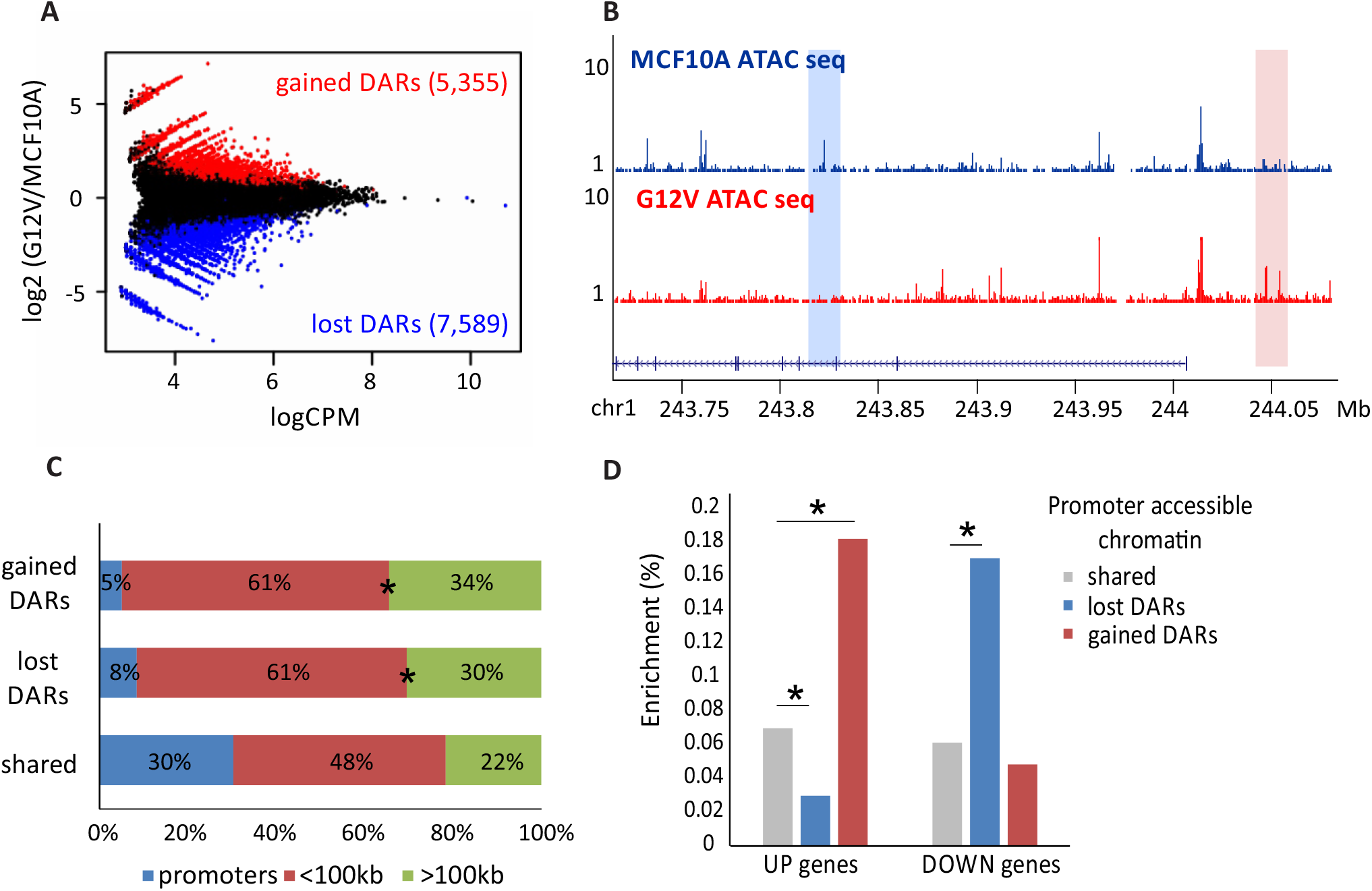
G12V HRas induced changes in the transcription regulatory landscape. **(A)** MA plot displaying the mean normalized counts (Counts Per Million (CPM), x-axis) versus the log 2 fold change between MCF10A and G12V MCF10A cells (y-axis), of the ATAC seq peaks. Red and blue points represent unique G12V (gained) and unique MCF10A (lost) peaks, respectively, as declared by MACS analysis. **(B)** Representative examples of lost (blue box) or gained (pink box) DARs. Chromosomal coordinates in Mb of human hg19 genome build are indicated in the bottom. **(C)** Distribution of regulatory sites relatively to genes: promoter proximal (+500bp, −1000bp from TSS, red), mid-range (+-100kb, blue) and far-range (green) in gained or lost DARs or accessible regions shared by both cell types. *p< 0.001, proportional test. **(D)** % of regulatory sites near promoters of up-regulated genes in G12V MCF10A cells (UP genes) or down-regulated genes in G12V MCF10A cells (DOWN genes) from total regulatory sites near promoters. *p< 0.001, proportional test.

The data points to enhancers as key elements in regulation of transcriptional reprogramming during oncogene-induced tumorigenesis in this model. Thus, in order to define regulatory pathways that are important in this process, it is essential to connect between DARs and their distant target genes. For this we used available high-resolution Hi-C data from multiple cell types that partitions the whole human genome to TADs - segments of high spatial associations ^23^. Using 4C, we first tested for a few differentially expressing and a few stably expressing genes, whether their Hi-C defined topological domains are stable between G12V MCF10A cells and their parental counterparts. As shown in Fig. 4A and Supplementary Fig. S1, indeed there is no change in the domains of the 5 genes we tested. We therefore took advantage of the comprehensive TAD database available ^23^ as a regulatory organizational framework. As shown in Fig. 4B, regulatory elements can be found close to a gene’s transcription start site, however not in the same topological domain, while a further away regulatory element is found in the same topological domain which is more biologically relevant. Thus using TADs as the spatial platform to connect between genes and their regulatory elements is a robust and biologically relevant approach.

**Fig. 4.**
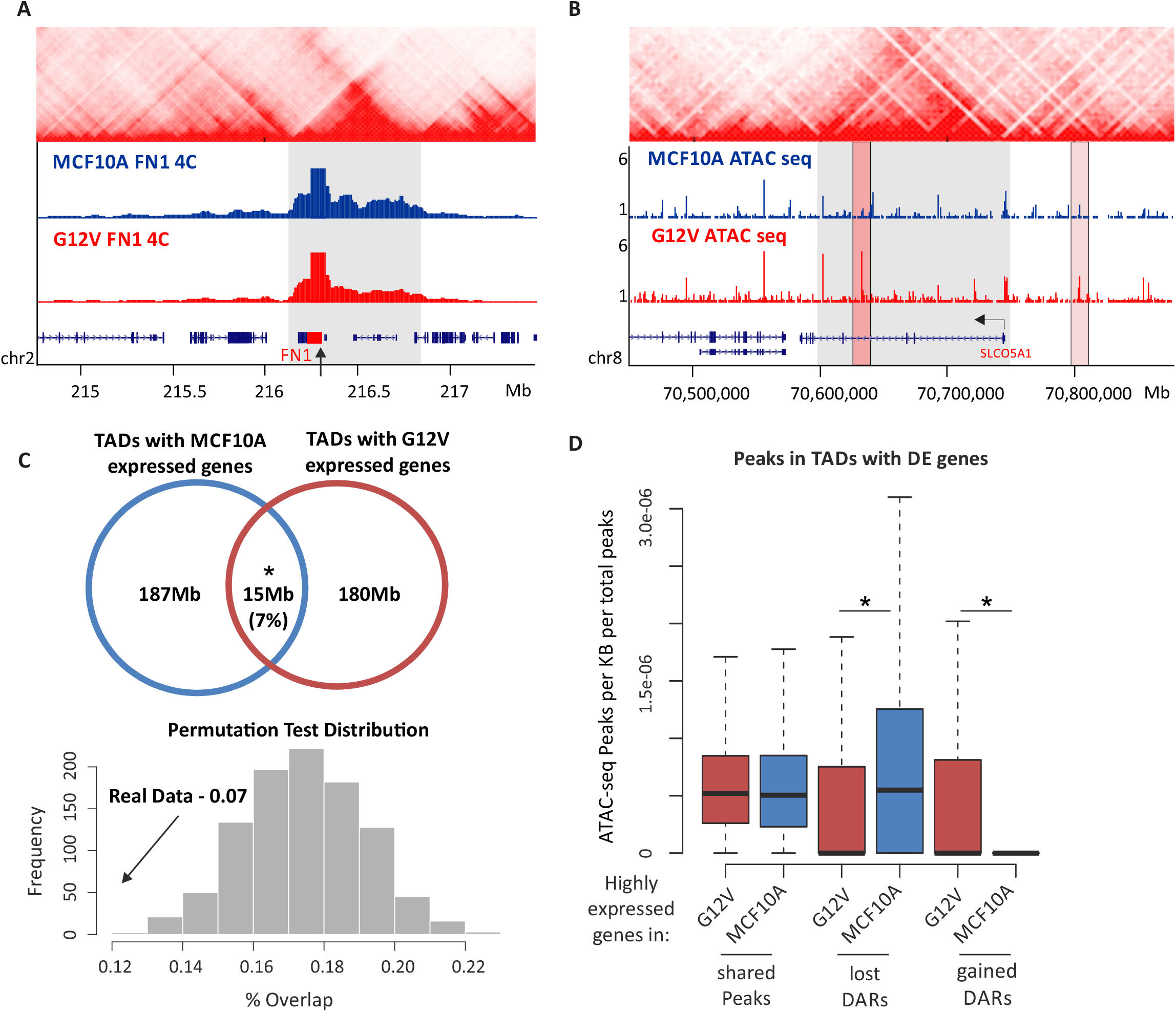
TADs as the spatial framework of transcriptional regulation. **(A)** Example of 4C-seq profile of the down-regulated FN1 gene (marked with black arrow) in MCF10A (blue track) and G12V MCF10A cells (red track) showing there are no changes in FN1 domain (marked with gray box) borders after transformation. Hi-C data from GM12878 is shown on the top ^23^. Chromosomal coordinates in Mb of human hg19 genome build are indicated in the bottom **(B)** Example for a domain of an up-regulated gene, SLCO5A1, (marked with gray box, TSS marked with black arrow) as defined from Hi-C data (shown on the top, ^23^) and the ATAC-seq data in MCF10A and G12V MCF10A cells. Right pink box indicates a unique regulatory site relatively close to the TSS which is outside of the TAD and the left pink box indicates a further away regulatory site that is within the TAD. Chromosomal coordinates in Mb of human hg19 genome build are indicated in the bottom. **(C)** Top - Venn diagram showing the overlap in bp between the domains of up-regulated genes and down-regulated genes after HRas transformation. *p<0.001, permutation test. Bottom-histogram showing permutation test results for the degree of overlap between up-regulated and down-regulated genes in the same TAD. Real overlap (shown with an arrow) is significantly lower than the overlap expected given random distribution. **(D)** Boxplots showing the number of peaks per kb in up (red) or down (blue) domains. Left – MCF10A regulatory sites, right-G12V regulatory sites. *p<0.01, Wilcox test.

The 3D organization of the genome has been shown to provide functional framework to transcriptional regulation. Significantly, in line with this notion, we found that up-regulated and down-regulated genes are segregated between TADs and fall together in the same TAD much more rarely than expected by chance (Fig. 4C). Therefore, our data supports that also in the case of oncogene-induced carcinogenic transformation, TADs are functionally relevant to transcriptional regulation, thus constitute an adequate framework to couple between transcriptional activity and regulatory loci as defined by chromatin accessibility. Importantly, cell-type specific (parental or G12V MCF10A) DARs are enriched within TADs containing cell-type specific expressed genes (Fig. 4D) suggesting that chromatin accessibility and genes are not only physically but also functionally linked within the 3D domains.

To discover candidate TFs regulating transcriptional reprogramming in this model of oncogene-induced transformation, we applied motif discovery analysis on different groups of regulatory sites defined by ATAC-seq, that are associated with differentially expressed genes in the same TAD.

Analysis of gained DARs, revealed enrichment for binding motifs of several TFs including ETV1 and Fli1 pro-oncogenes of the ETS family of transcription factors. Increased activity of ETS transcription factors was shown to be involved in all stages of tumorigenesis of several solid tumors, including prostate and breast cancer ^24,25^. RNA-seq analysis uncovered several ETS factors which are differentially expressed in G12V cells and may be involved in transcriptional reprogramming. One of the motifs that was found enriched in gained DARs is the CTCF motif. Interestingly, the CTCF motif is enriched in gained DARs located in TADs of both up-regulated and down-regulated genes in G12V cells (Fig. 5A). This suggests that the enrichment of the CTCF motif is associated with gained regulatory activity upon HRas oncogenic activation that is associated with both gene activation and gene repression. However, according to RNA-seq data the levels of CTCF did not differ between G12V and parental MCF10A cells (Fig. 5B). Moreover, CTCF protein levels, as tested by western blot analysis did not change either (Fig. 5B). To validate that indeed there is a change in CTCF binding upon activation of the G12V HRas oncogene, we performed chromatin immunoprecipitation coupled with next generation sequencing (ChIP-seq) for CTCF in G12V MCF10A cells and their parental counterparts. This analysis revealed 30,642 ChIP-seq peaks in G12V MCF10A cells and 22,376 peaks in the parental MCF10A cells. As was predicted from the motif analysis, CTCF ChIP-seq data shows significantly higher CTCF binding in gained DARs compared to lost DARs located in TADs of both up-regulated and down-regulated genes (Fig. 5C, D). This strongly suggests the involvement of CTCF in the regulation of transcriptional reprogramming upon HRas transformation in this system. Interestingly, this occurs via redistribution and probably increase of genomic CTCF binding upon HRas oncogene-induced transformation without change in the level of CTCF expression.

**Fig. 5.**
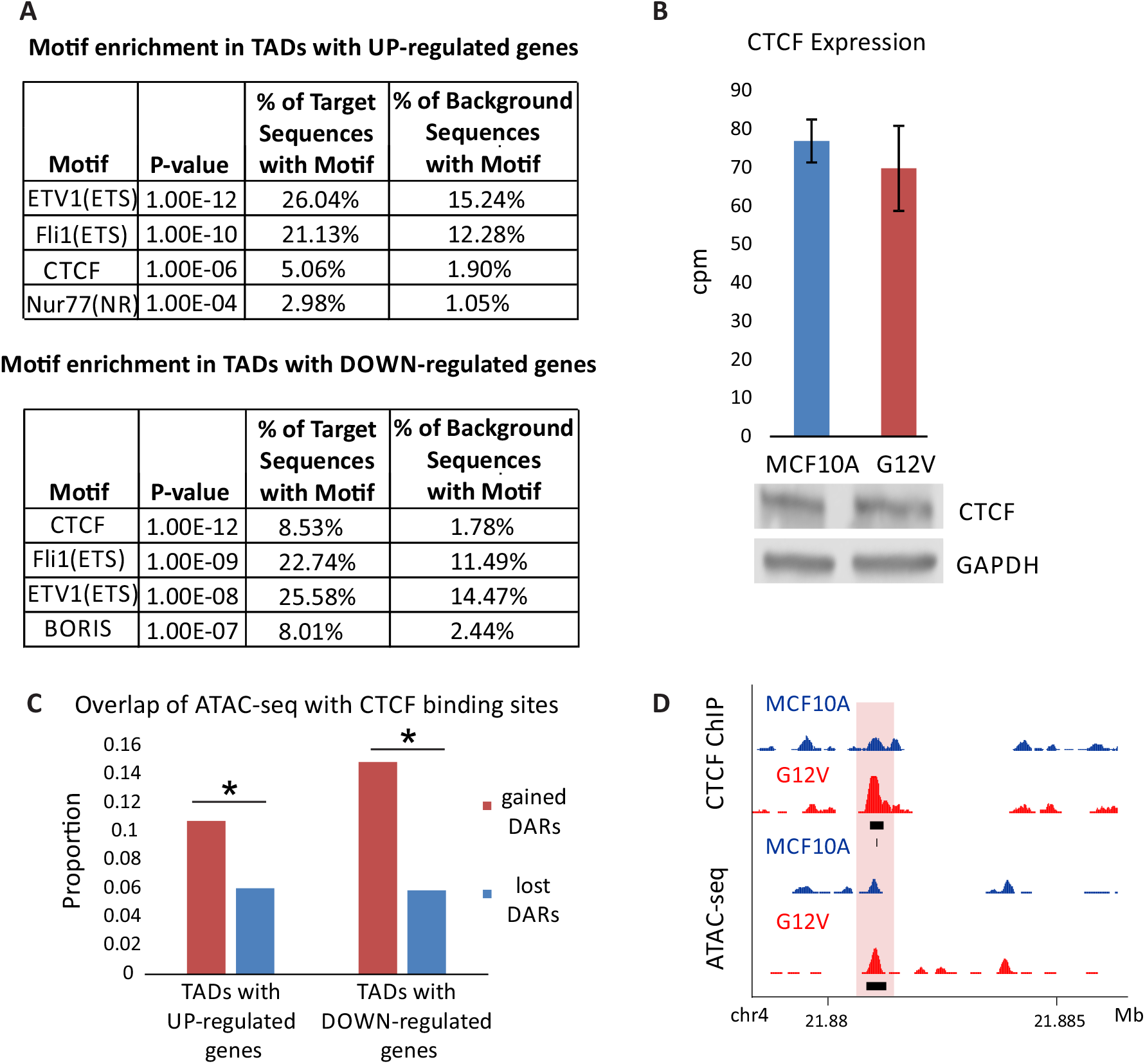
CTCF motif and binding sites are enriched in gained DARs. **(A)** Top motifs enriched in gained DARs within up TADs (upper table) or down TADs (lower table). Background group is the lost DARs. **(B)** Expression levels of CTCF in MCF10A (blue) and G12V (red) cells determined by RNA seq (top). Error bars represent standard deviation of two biological replicas. CTCF protein levels in MCF10A and G12V MCF10A cells as measured by western blot analysis (bottom). Representative result of at least two biological repeats is shown. **(C)** % of gained (blue) and lost (red) DARs overlapping CTCF binding sites in specific up and down TADs. *p<0.001, proportional test. **(D)** Example for a gained DAR that overlaps with gained CTCF binding site (pink box). Black boxes indicate peaks, the black line indicates CTCF motif. Chromosomal coordinates in Mb of human hg19 genome build are indicated in the bottom.

In lost DARs associated with differentially expressed genes, the most significantly enriched motif is the binding motif of p53 (Fig. 6A). This is in agreement with the fact that p53 is one of the top enriched upstream regulators with multiple of its targets being differentially expressed following HRas G12V overexpression in MCF10A cells (Fig. 2E). Since the highest enrichment of p53 motif was in lost DARs associated with genes that were down-regulated following G12V HRas transformation, we sought to confirm that such genes are indeed regulated by p53 in the parental MCF10A cells. To measure the likely direct transcriptional response to p53, RNA was extracted following a 4-hour Nutlin-3a treatment and sequenced. Gene expression changes were overall moderate. Importantly genes that were down-regulated following HRas induced transformation and were located within TADs harboring lost DARs that contain p53 motif showed the highest and significant response to p53 activation (Fig. 6B, and Supplementary Fig. S3). We therefore checked the level of p53 for changes that could explain the mis-regulation of its target genes. Surprisingly *TP53* transcript and p53 protein levels, as well as its sub-cellular distribution remain stable between G12V and MCF10A cells (Fig. 6C, D). Moreover, we confirmed that *TP53* gene is not mutated in the G12V HRas expressing cells (data not shown). Thus the reduction in accessibility at putative p53 binding sites and mis-regulation of p53 target genes in G12V MCF10A cells are not due to its downregulation or change in cellular localization. To assess directly whether indeed the landscape of p53 chromatin binding was altered in MCF10A and G12V cells, ChIP-seq was performed. Overall, 3,260 binding sites of p53 were found in both cell types, with 465 and 565 sites specific for parental MCF10A and G12V cells, respectively. As was predicted by the motif analysis, p53 binding is significantly higher in lost DARs relatively to gained DARs located in TADs of up-and down-regulated genes, reinforcing the direct role of p53 in transcriptional reprogramming in G12V cells (Fig. 6E). Notably, the variation in p53 binding was accompanied by significant decrease in the proportion of canonical p53 binding motif (from 93% to 69%, p < 2.2e-16, proportional test, in cell type-specific peaks (Supplementary Fig. S4), suggesting that p53 binding in G12V cells is mediated by another transcription factor. Interestingly, the binding motif of AP-1 family of transcription factors is highly enriched in G12V-specific p53 binding loci (24%) relatively to the entire ChIP-seq dataset in these cells (14%) and particularly relatively to the MCF10A-specific binding loci (6%), which is close to the background level. This supports a major role for p53 in the transcriptional reprogramming during the early stages of HRas oncogene induced transformation. Importantly, this reprogramming by p53 does not occur through changes in protein level or localization, but rather through changes in its chromatin binding.

**Fig. 6.**
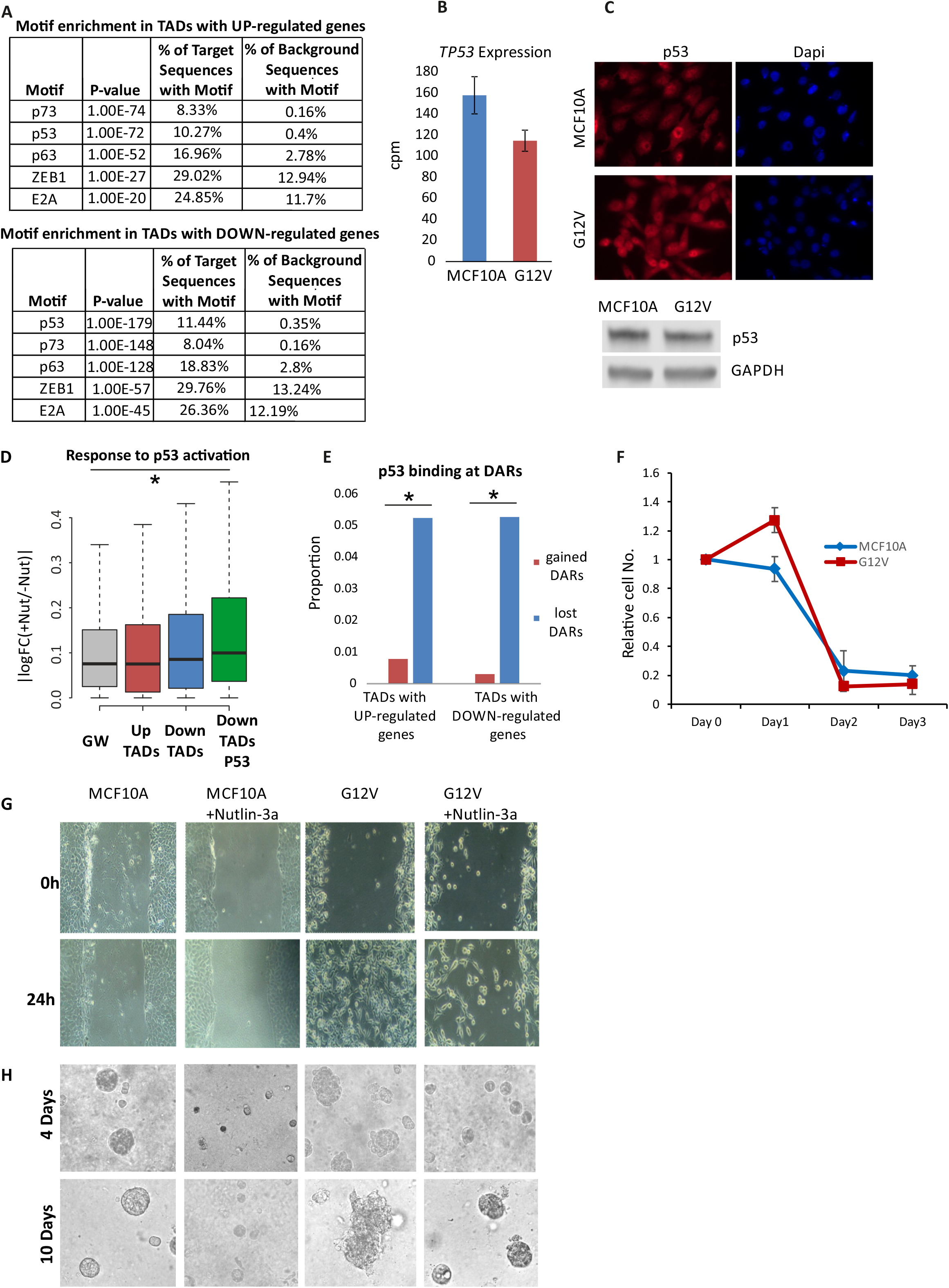
Redistribution of p53 binding underlie genetic reprogramming and caner phenotypes. **(A)** Top motifs enriched in unique Parental regulatory sites within up TADs (upper table) or down TADs (lower table). Background group is gained DARs. **(B)** RNA levels of *TP53* in MCF10A (blue) and G12V (red) cells determined by RNA-seq. Error bars represent standard deviation of two biological replicas. **(C)** Immunostaining of p53 (top panel) in MCF10A and G12V cells. p53 protein levels in MCF10A and G12V MCF10A cells as measured by western blot analysis (bottom). Representative results of at least two biological repeats are shown. **(D)** Variation in RNA levels |log2FC| following 4h Nutlin-3a treatment in MCF10A cells. Gray – all expressed genes, red –up-regulated genes after HRas transformation, blue – down-regulated genes after HRas transformation and green – down-regulated genes that have lost regulatory sites with a p53 motif in their domain. *p<0.001, Wilcox test. **(E)** % of gained (red) and lost (blue) DARs overlapping p53 binding sites in specific up and down TADs. *p<0.001, proportional test. **(F)** Cell proliferation with 5μM Nutlin-3a was measured by XTT assay. The relative number of cells, compared to day 0 is presented for MCF10A (blue line) and G12V MCF10A cells (red line). **(G, H)** Representative images showing the migration capability and growth pattern of MCF10A and G12V cells with or without Nutlin-3a.

Finally, we asked to what extent the cancerous phenotypes of the transformed cells are related to p53. While activation of p53 by Nutllin inhibited cell proliferation of both, parental and transformed cells, this response was initially attenuated in G12V cells, which also maintained their capacity to migrate (Fig. @@@). The disorganized growth of G12V cells in 3D cell culture setup is a hallmark of cell transformation (Fig.). p53 activation by mild Nutlin-3a treatment reversed the effect of HRAS overexpression on mammosphere formation, while reduced the size of the mammospheres from MCF10A cells. These results indicate that the redistribution of p53 binding did not eliminate its protective capacity but may have diminished some of its arms.

## Discussion

Cancer development is associated with altered gene expression programs which are key in the acquisition of biological capabilities that drive tumorigenesis. Defining the regulatory networks underlying carcinogenesis associated transcriptional reprogramming holds great promise for identifying key transcription factors in this process that may not harbor mutations or change expression patterns. Transforming cells by over expressing oncogenes is widely used and has been instrumental in understanding molecular mechanisms involved in malignancy ^26^. These cell-based models allow to explore processes occurring in early stages of oncogene-induced transformation. Using a cell-based mammary epithelial model we find that transformation induced by overexpression of oncogenic HRas is associated with dramatic changes in gene expression. Indeed, the changes we report are characteristic of a carcinogenic transformation process.

Regulatory sites are known to malfunction and thus cause major gene expression alterations related to cancer ^27,28^. Several chromatin characteristics are used as proxy for activity of regulatory elements, including DNA methylation status, nucleosome occupancy, specific histone species and modifications in flanking regions. These were used in several studies to demonstrate major alterations in the DNA regulatory landscape in various cancer cell lines and tumor cells from different cancer types ^28–32^. By assaying chromatin accessibility, we demonstrate that even at early stages of transformation, induced by over expression of oncogenic G12V HRas variant, major alterations in the regulatory landscape are evident. Similar modulation of the regulatory landscape was described previously during Ras-dependent oncogenesis in drosophila ^33^ and using H3K27ac marking following disruption of the ERK signaling pathway in MEF cells ^34^. Strikingly, examination of the distribution of regulatory sites between promoters and enhancers revealed that enhancers are dramatically enriched in the varying fraction of active regulatory elements between transformed and non-transformed cells, illustrating that they have a substantial role in transcriptional reprogramming in oncogenic HRas induced transformation.

In order to interrogate more thoroughly specific transcriptional regulation of differentially expressed genes, it is necessary to assign altered regulatory elements to their target differentially expressed genes. Distance between regulatory elements and their target genes was shown to vary in the range of hundreds of kbs ^15^, however, regulation of gene expression is constrained by the 3D organization of the genome ^35^. TADs are topological domains in the length of hundreds of kbs of high frequency physical association which are considered stable across different cell types and to some extent even different species ^23,36,37^. Therefore TADs are considered a framework within which enhancer-promoter dynamic interactions occur ^16,35^. Importantly, TADs were shown to act as co-regulated functional units in different processes from long-term differentiation processes ^37^ to short-term responses to external signals ^20,21^. Strikingly also in the process of oncogene-induced transformation we find strong segregation between up-regulated and down-regulated genes among TADs, which strongly supports that TADs still comprise the regulatory context within which transcriptional reprogramming occurs during early stages of transformation. Moreover, we confirm that there are no major changes in TAD boundaries for a number of differentially expressed genes as a results of the transformation process.

Motif enrichment in gained DARs that are associated with differentially expressed genes can infer transcriptional regulation activity in transformation-associated transcriptional reprogramming. The transcriptional regulator CTCF was found to be enriched in specific active regulatory sites in G12V MCF10A cells despite lack of change in its expression. Mutations in CTCF are frequently found in breast tumors ^3,38^, some of which have been shown to affect its DNA binding. Interestingly, the effect of mutations on CTCF binding was found to be non-uniform, for instance different mutations in its zinc finger 3 domain led to selective inhibition of binding to different targets suggesting that there is a tumor-specific change of function rather than loss of function ^38^. Furthermore, CTCF was shown to be elevated in breast cancer cell lines and breast tumors ^39^, its overexpression was suggested to protect tumor cells from induction of apoptotic cell death ^40^ while its downregulation in breast cancer cell lines was shown to be associated with reduced cell proliferation ^41,42^, both effects via transcriptional regulation of target genes. In line with these results we find overall increased chromatin binding of CTCF in G12V MCF10A cells, although without detectable elevation in its gene expression levels.

P53 is a known tumor suppressor and the TP53 gene is frequently mutated in breast cancer ^3^. We found the p53 binding motif to be strongly enriched in domains of differentially expressed genes in response to HRas oncogenic activity. Moreover, these genes were indeed regulated by p53 as activation of p53 by Nutlin specifically increased their expression level. However, like CTCF, we found that p53 chromatin binding is redistributed while its expression levels are not changed as well as its localization in the cell. The involvement of p53 in cancers is associated with its DNA binding and transcriptional regulation activity ^43^. Our results suggest that distinct genomic p53 binding patterns reported in cancer cells ^44–46^ may be important for the development of capabilities during the progression of cancer. Given p53 cancer-specific binding profile it is unlikely that p53 alone reprogram the chromatin landscape^47^. Interestingly, redistribution of p53 binding is accompanied with decrease in the proportion of its binding motif and increase in AP-1 binding motif. This suggests a possible role for members of the AP-1 family of transcription factors in modulating p53 activity in the transformed cells by direct association with p53 binding sites ^48^. Functionally, this modulation may relate to transcriptional suppression as p53 represses target genes of TFF2 via the AP-1 motif ^49^.

The 3D conformation of the genome is the framework within which transcriptional regulation takes place. Here we take advantage of this framework together with DNA accessibility as proxy for active regulatory elements, to infer from changes in gene expression upon HRas oncogene over expression in normal breast epithelial cells, on regulatory networks that are dis-functioning at the first stages of oncogene induced carcinogenesis. Chromosome structure Hi-C data allowed combining regulatory loci that change their activity with their distant transcriptionally responsive gene-targets in their biologically relevant 3D context on a genome-wide scale. This focused approach coupled with motif discovery analysis revealed two key regulatory factors, namely CTCF and p53, that regulate the transcriptional reprogramming associated with HRas oncogenic cellular transformation. Noteworthy, these factors carry out this transcriptional reprogramming, while their RNA, protein levels and sub-cellular localization remain invariable, therefore could have not been identified by differential expression analysis. Our ATAC-seq and ChIP-seq data support that this effect is occurring through changes in DNA binding patterns. Thus combination of differential expression analysis, and DNA accessibility using the framework of the 3D organization of the genome is a powerful tool to identify in an unbiased manner key regulatory pathways that orchestrate transcriptional reprogramming during early stages of cancer development.

## Acknowledgments

This work is supported by the Israel Science Foundation (grant 748/14), Marie Curie Integration grant (CIG)-FP7-PEOPLE-20013-CIG-618763 and I-CORE Program of the Planning and Budgeting Committee and The Israel Science Foundation grant no. 41/11. We thank the BIU imaging facility for helping with microscopy.

## Author contributions

M.S., B.Z.A., M.T., O.L., A.C. and Z.S. performed experiments, M.S., A.S.P., T.K., Z.S. and O.H. analyzed data. M.S., A.S.P., Z.S. and O.H. designed experiments. M.S., A.S.P., Z.S. and O.H. wrote the manuscript.

## Methods

### Cells and treatments

MCF10A cells were grown as previously described (Debnath et al.) in DMEM media (Biological Industries) supplemented with 100 ng/ml cholera toxin (Sigma), 20 ng/ml epidermal growth factor (EGF, Peprotech), 0.01 mg/ml insulin (Sigma), 500 ng/ml hydrocortisone (Sigma), 1% penicillin-streptomycin (Biological Industries), 5% horse serum (Biological Industries). Introduction of the G12V HRas oncogene was done via lentiviral transduction along with a GFP expression vector in order to track transduction efficiency. 72 hours post lentiviral transduction hygromycin was added to the media in order to select for cells carrying the G12V HRas expression vector. Nutlin treatment was done by adding 10µM Nutlin-3A (Sigma) to the media for the periods of time indicted.

### Lentiviruses

Lentiviral particles were prepared by cotransfecting either the G12V HRas over expression vector pWZL hygro HRas V12 (addgene #18749) or a GFP expression vector with packaging vectors (CMVΔR8.91, CMV-VSV-G) into 293T cells using Mirus TransLTi (Mirus Bio LLC, Madison, WI) according to the manufacturer’s instructions. Medium containing viral particles was collected 2 and 3 days post transfection.

### Proliferation assay

Cell proliferation was measured for 3days using an XTT based cell proliferation kit (Biological industries), according to the manufacturer’s instructions.

### Colony Formation assay

100 cells per well were seeded in 6 well-plates in triplicates. After two weeks, cells were fixed and stained with Giemsa stain. Colonies larger than 5mm were counted.

### Soft agar assay

2×10^4^ cells were seeded in MCF10A standard media containing 0.3% 2-Hydroxyethyl Agarose on top of a solidified layer of MCF10A media containing 0.6% 2-Hydroxyethyl Agarose. Plates were incubated until colonies were visible by naked eye at which point they were counted.

### Anchorage independent cell death assay

5×10^3^ cells were plated onto poly-HEMA coated 12-well plates to prohibit attachment. After 4 days in suspension, cells were collected from the wells and live cells were counted manually using Trypan blue exclusion.

### Matrigel invasion assay

Cells were placed in the upper chamber of a Boyden migration chamber. The upper and lower chambers were separated by matrigel coated PVP-free, 8-mm pore size polycarbonate filters (Costar Scientific). EGF containing conditioned medium of 3T3 fibroblasts was placed in the lower chamber. After an overnight incubation the filters were fixed and stained with Diff-Quick System (Dade Behring, Inc.) and cells on the lower surface were counted. Each assay was done in triplicates.

### RNA extraction, cDNA synthesis and qPCR

Total RNA was extracted from the cultured cells using Quick-RNA MiniPrep kit (ZYMO RESEARCH) following manufacturer’s instructions, reverse transcribed using Quanta Bioscience qScript cDNA synthesis kit (95047-100) following manufacturer’s instructions. cDNA was measured by real time PCR (Bio-Rad S1000) using sybr green mix (Bio-Rad) with primers spanning exon-intron junctions (Supplementary Table S3) and normalized to GAPDH transcript. Results show average and SD of three replicates.

### Wound healing assay

25×10^4^ cells/well were seeded in 12 well plate wells to form a 100% confluent layer. One day after the layer was wounded using the 10μl pipet tip and monitored for wound healing at 0 h and 24h in minimal growth factor medium.

### 3D culture assay

Matrigel was incubated on ice overnight, then 3000 cells were seeded on a solidified layer of growth factor reduced Matrigel measuring approximately 1–2 mm in thickness. The cells were grown in an assay medium containing 5 ng/ml EGF and 2% Matrigel then incubated in CO2 incubator at 37 °C.

### RNA-seq

Total RNA was extracted from cells using RNA purification kit (GeneAll) according to the manufacturer’s instructions. RNA quality was measured on a Bioanalyzer (Agilent) and only RNA with RIN score >9 was used for library preparation. Messenger RNA (mRNA) was enriched from 1 μg of total RNA by Poly(A) mRNA Magnetic Isolation Module (New England Biolabs) according to the manufacturer’s instructions. cDNA libraries were constructed using the NEBNext Ultra RNA Library Prep Kit (New England Biolabs) following the manufacturer’s protocol. Library concentration was measured by DNA High Sensitivity Kit (Invitrogen) on a Qubit fluorometer (Invitrogen). Library quality and fragment sizes were assessed on a Bioanalyzer (Agilent) or on a Tape station (Agilent). RNA-Seq libraries from at least two biological replicas for each condition were sequenced on Illumina Hi-seq 2000 platform.

### ATAC-seq

ATAC-seq was performed as previously described (Buenrostro 2013). Briefly, cells were lysed and nuclei counted. Transposition reaction was performed on 10^5^ nuclei using 5μl of Nextera TDEI enzyme (Illumina) for 30 minutes at 37°C. Library was amplified using NEBNext High-Fidelity 2× PCR Master Mix (New England Biolabs). Library concentration was measured by DNA High Sensitivity Kit (Invitrogen) on a Qubit fluorometer (Invitrogen). Library quality and fragment sizes were assessed on a Bioanalyzer (Agilent). ATAC-Seq libraries from two biological replicas for each condition were sequenced on Illumina Hi-seq 2000 platform.

### ChIP-seq

Cells were cross-linked for 10 min at 37°C in 1% formaldehyde followed by quenching with 125 mM glycine for 10 min. for CTCF, crosslinked cells were first lysed in cell lysis buffer (10mM Hepes pH7.5, 85mM KCl, 1mM EDTA, 1% NP-40) supplemented with protease inhibitors, resuspended in RIPA buffer (10 mM TrisHCl, pH 7.6, 1 mM EDTA, 0.1% SDS, 0.1% NaDeoxycholate, 1% Triton X100) supplemented with protease inhibitors, and sonicated for 40 cycles of 30s ON and 30s OFF (Bioruptor sonicator, Diagenode). Cleared chromatin was incubated overnight with 10 μg α-CTCF (Millipore 07-729) or 5 μg α-p53 (DO-1, Santa-Cruz) and additional 2 hours with 40ul protein A/G magnetic beads (ChIP grade, Pierce). Complexes were washed twice with RIPA buffer, twice with RIPA buffer supplemented with 300mM NaCl, twice with LiCl buffer (10mM Tris 7.5, 1mM EDTA, 0.25M LiCl, 0.5% NP40, 0.5% Na-Deoxycholate), once with TE buffer (10 mM Tris, 1 mM EDTA, pH8) supplemented with 0.2% triton and once in TE buffer. For p53, cells were treated with 10uM Nutlin-3a (Sigma) for 4 hrs before fixation. Fixed cells were lysed with SDS Lysis Buffer (1% SDS, 50mM Tris pH 8.1, 10mM EDTA) supplemented with protease inhibitor and sonicated for 560 seconds (ME220 sonicator, Covaris). Cleared chromatin was diluted 1:10 with dilution Buffer (16.7mM Tris-HCl 8.1; 1.2mM EDTA; 167mM NaCl; 1.1% Triton) and incubated over night with 5 μg α-p53 (DO-1, Santa-Cruz) bound to magnetic beads (Dynabeads Protein A). Chromatin was washed with low salt buffer (20mM Tris-HCl 8.1; 2mM EDTA; 150mM NaCl; 1% triton; 0.1% SDS), high salt buffer (20mM Tris-HCl 8.1; 2mM EDTA; 500mM NaCl; 1% triton; 0.1% SDS), LiCl buffer (10mM Tris-HCl 8.1; 1mM EDTA; 1% NP-40; 250mM LiCl) at 4°C, and twice with TE (10mM Tris-HCl 8.1; 1mM EDTA) at room temp. Complexes were eluted with Elution buffer (10mM Tris-HCl 8.1; 1mM EDTA; 200mM NaCl; 1% SDS). For both antibodies, crosslinks were reversed with 1 mg/mL Proteinase K overnight at 65°C. Purified DNA was used to prepare sequencing libraries using NEBNext UltraII DNA Library Prep Kit (New England Biolabs). Library concentration was measured by DNA High Sensitivity Kit (Invitrogen) on a Qubit fluorometer (Invitrogen). Library quality and fragment sizes were assessed on a Bioanalyzer (Agilent). ChIP-seq libraries from two biological replicas for each condition were sequenced on Illumina Hi-seq 2000 platform.

### 4C-seq

4C was performed as previously described (Schwartz, biotechniques, Hakim, Genome Res 2012). Proximity ligation junctions, reflecting in vivo spatial proximity, were generated with HindIII (New England Biolabs), followed by circularization with Csp6I (Thermo Scientific). Chromosomal contacts with the baits were amplified with inverse PCR primers (Supplementary Table S2) using Platinum Taq DNA Polymerase (LifeTechnologies). 4C-seq libraries were sequenced on Illumina Hi-seq 2000 platform.

### Immunostaining

Cells were fixed with 3.7% formaldehyde, permeabilized with 0.1% triton and blocked with 2% BSA. p53 was detected using a mouse monoclonal antibody (SC-126, Santa Cruz Biotechnology) followed by a Cy3 conjugated anti-mouse antibody (Jackson Immunoresearch Laboratories). Cells were stained with Dapi and viewed on an Axioimager fluorescent microscope (Zeiss).

### Western blot analysis

Cells were lysed directly in sample buffer and protein extracts were separated on a 4-20% polyacrylamide gel (Bio-Rad). Blotted nitrocellulose membranes were incubated with primary antibodies (mouse -p53, SC-126, Santa Cruz Biotechnology; rabbit -CTCF, Millipore 07-729; rabbit -GAPDH, cst-2118, Cell Signaling Technology) followed by detection with HRP conjugated secondary antibodies (Jackson Immunoresearch Laboratories). ECL was performed using EZ-ECL kit (Biological Industries) and imaged on an ImageQuant chemiluminescence camera (GE Healthcare).

### RNA seq analysis

Alignment was done using TopHat ^52^. Reads count on transcripts was done using HTSeq ^53^. Expression and differential analysis was done using ‘edgeR’ ^54^ that assigns to each gene a False Discovery Rate (FDR) value and calculates the log Fold Change (logFC) between the two conditions. Genes with FDR <0.05 and logFC greater than |0.5| were considered differentially expressed.

Functional enrichment was done using the IPA (Qiagen, ^55^).

### ATAC seq and ChIP-seq analysis

Alignment was done using Bowtie ^56^, allowing only unique reads to be considered. Since the replicas were well correlated (Supplementary Fig. S2), reads from both replicas of each condition were combined for further analysis. For detecting regions of local read enrichment (peaks), MACS algorithm ^57^ was applied with default parameters for ChIP-seq. For ATAC-seq, read start site was adjusted to represent the center of the transposon binding event as described ^58^ and the following parameters for MACS were applied – --tsize=51 --nomodel --shiftsize=75 --llocal=25000 -p 1e-04. Differentially accessible loci between MCF10A and G12V cells were detected by MACS algorithm, using one sample as the background of the other. Only peaks that were both detected by MACS analysis in either sample and in the comparative MACS analysis were considered as unique peaks and were used for further analysis.

For calculating the distribution of ATAC-seq peaks relatively to gene features, refseq annotations were used for TSS coordinates of each gene, and the longest transcript as gene body. Overlap of at least 1 bp between ATAC-seq peak with gene promoters (1000bp upstream and 500bp downstream to the TSS) or with distant regulatory regions (100kb upstream and downstream to the TSS of each gene) was counted. Regulatory sites that did not overlap one of the two features were considered as far regulatory sites. Overlap analysis was performed by intersectBed from Bedtools ^59^. Each gene or peak was counted once.

### 4C-seq analysis

Reads were sorted, according to their barcodes, to different fastq files for each bait and condition and aligned to the human (hg19) genome using BOWTIE ^56^. Reads were then counted for each HindIII site. For domains detection the number of reads on each HindIII site was counted in sliding windows of 50Kb with 25Kb steps. Hi-C heatmaps are from the 3D genome browser ^60^.

### Association of genes and TADs

For genome-wide assignment of gene to topologically associated domains (TADs), TADs from high resolution Hi-C data of 7 human cell lines ^23^ were merged to one dataset and the smallest overlapping TAD was assigned for each gene. “Up” or “down” TADs were declared according to their containing up-or down-regulated genes after transformation by G12V HRas. Overlap between TADs, regulatory sites and genes was calculated by intersectBed from Bedtools ^59^.

### Motif discovery analysis

To uncover DNA binding motifs of possible transcription factors (TFs) which are associated with regulatory loci, HOMER ‘findMotifsGenome.pl’ program ^61^ with default parameters was applied for a 150bp window with the highest read count within each peak (summit) that was defined using custom made script in R. The enrichment of motifs in regulatory sites from MCF10A was measured against G12V cells and vice versa.

### Data availability

The sequencing data has been deposited to the GEO database (accession number GSEXXXX).

