## Supplemental data for "Genomic Retargeting of Tumor Suppressors p53 and CTCF Promotes Oncogenesis"

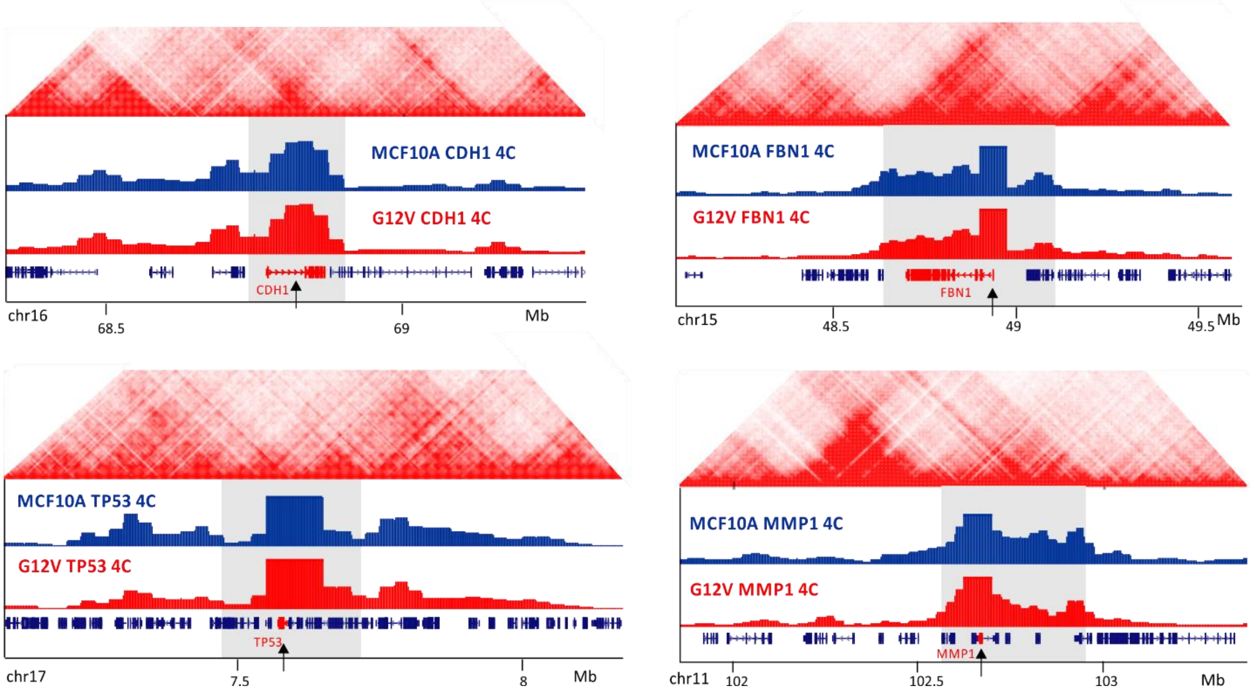


**Supplementary Figure S1. 4C profiles.** Four examples of 4C-seq profiles: CDH1 (down-regulated gene), FBN1 (up-regulated gene), TP53 (unchanged gene) and MMP1 (low expressing gene), in MCF10A and G12V MCF10A cells. Domains are marked with gray box, genes marked with black arrow. Hi-C data from GM12878 is shown on the top [23]. Chromosomal coordinates in Mb of human hg19 genome build are indicated in the bottom.

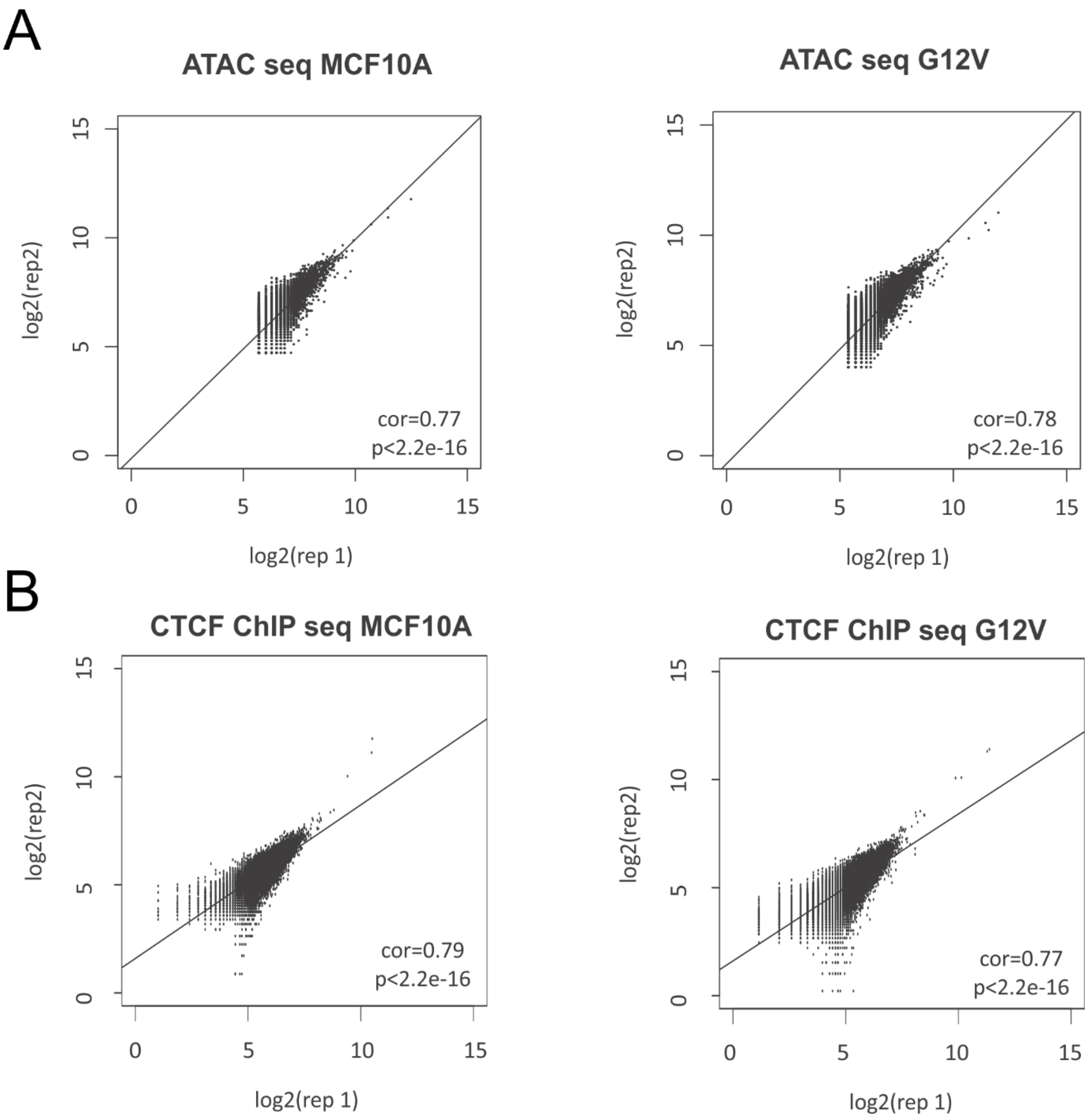


**Supplementary Figure S2. ATAC-seq and ChIP-seq correlations**. Correlation plots between ATAC-seq replicas (A) and ChIP-seq replicas (B). log2CPM of replica 1 within peaks was plotted against the log2CPM of replica 2 within peaks. Values for Pearson correlation and test are shown.

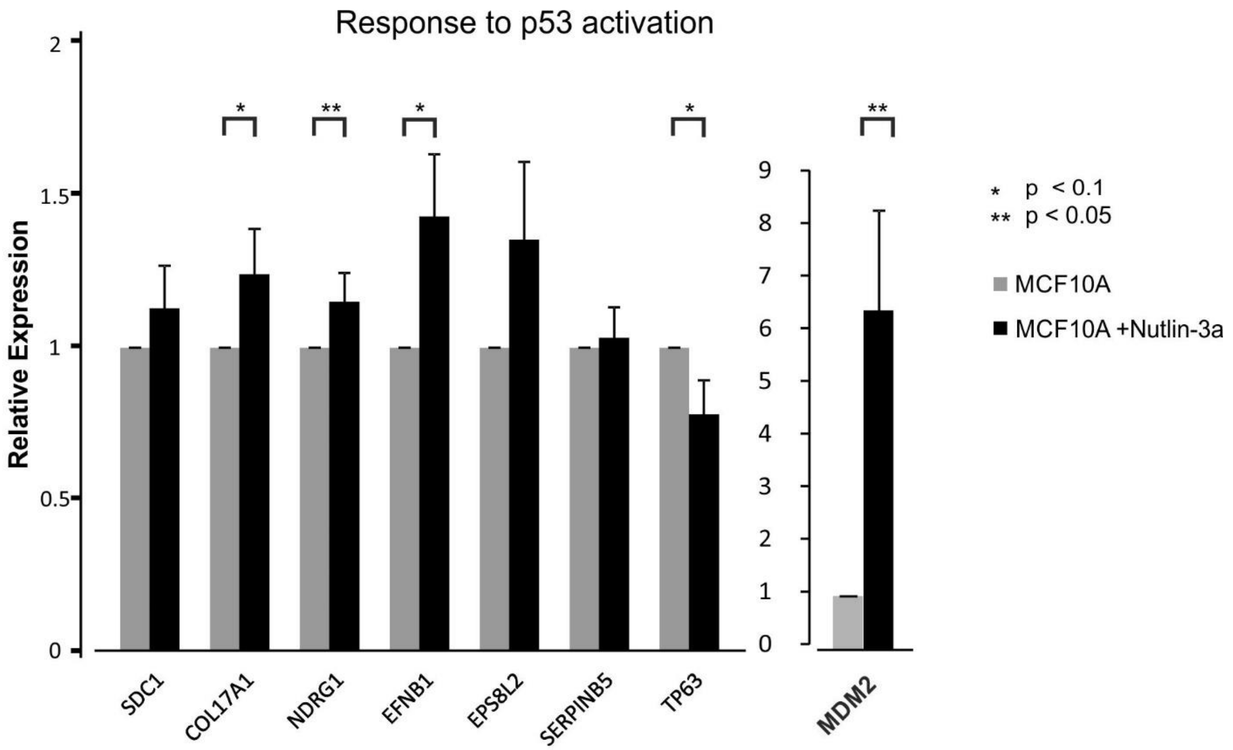


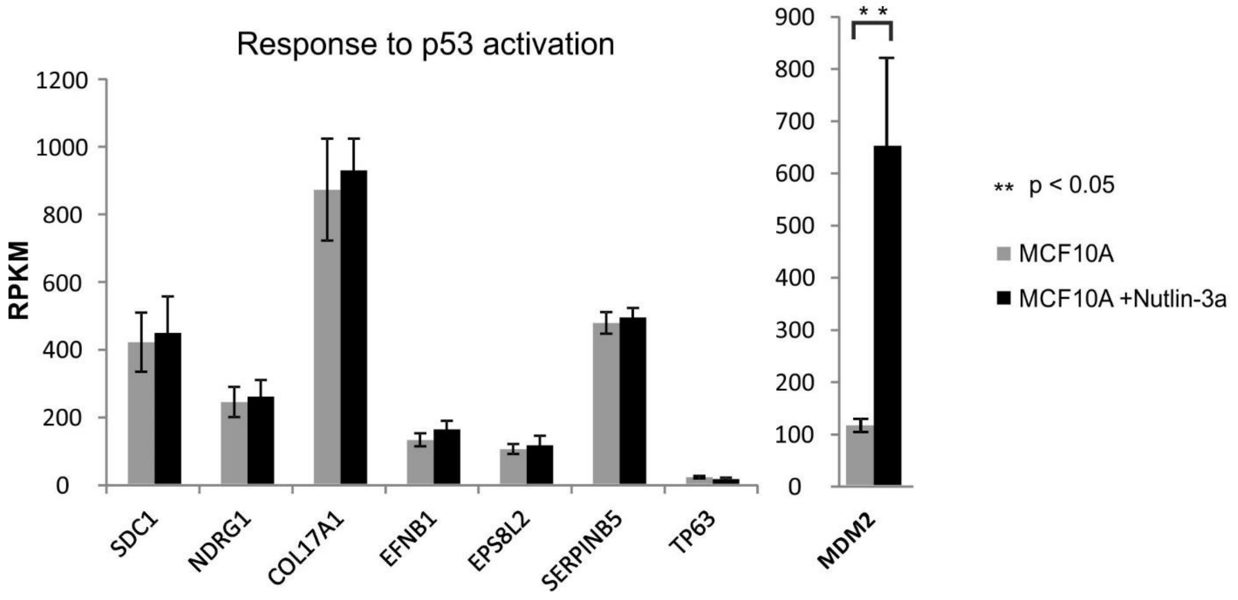


**Supplementary Figure S3. Transcriptional response to p53 activation** by Nutlin3a in MCF10A cells measured by real time PCR (top) and RNA-seq (bottom). genes in this figure were down regulated after HRas-induced transformation and contain p53 motif in their lost DARs. p-students t test. SD is presented.

### Motif enrichment in G12V-specific p53 ChIP peaks

Total Target Sequences = 465, Total Background Sequences = 48020

| Motif | Name | P-value | log P-pvalue | q-value (Benjamini) | # Target Sequences with Motif | % of Targets Sequences with Motif | # Background Sequences with Motif | % of Background Sequences with Motif |
|---|---|---|---|---|---|---|---|---|
| 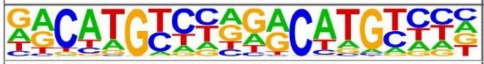 | p53(p53)/Saos-p53-ChIP-Seq(GSE15780)/Homer | 1e-478 | -1.102e+03 | 0.0000 | 320.0 | 68.82% | 485.0 | 1.01% |
| 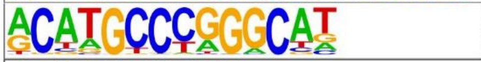 | p53(p53)/mES-cMyc-ChIP-Seq(GSE11431)/Homer | 1e-106 | -2.456e+02 | 0.0000 | 81.0 | 17.42% | 134.2 | 0.28% |
| 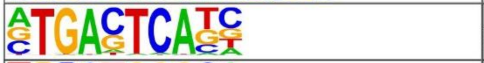 | AP-1(bZIP)/ThioMac-PU.1-ChIP-Seq(GSE21512)/Homer | 1e-38 | -8.939e+01 | 0.0000 | 111.0 | 23.87% | 2609.3 | 5.41% |
| 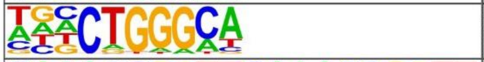 | ZNF416(Zf)/HEK293-ZNF416.GFP-ChIP-Seq(GSE58341)/Homer | 1e-23 | -5.492e+01 | 0.0000 | 196.0 | 42.15% | 10134.5 | 21.01% |
| 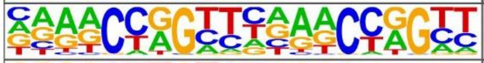 | Tcfcp2l1(CP2)/mES-Tcfcp2l1-ChIP-Seq(GSE11431)/Homer | 1e-17 | -4.133e+01 | 0.0000 | 37.0 | 7.96% | 579.2 | 1.20% |
| 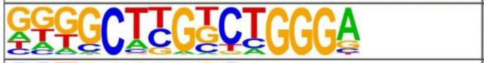 | Zfp809(Zf)/ES-Zfp809-ChIP-Seq(GSE70799)/Homer | 1e-15 | -3.632e+01 | 0.0000 | 57.0 | 12.26% | 1624.9 | 3.37% |
| 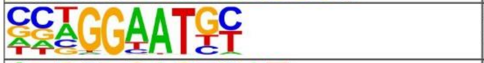 | TEAD4(TEA)/Tropoblast-Tead4-ChIP-Seq(GSE37350)/Homer | 1e-11 | -2.723e+01 | 0.0000 | 83.0 | 17.85% | 3735.6 | 7.74% |
| 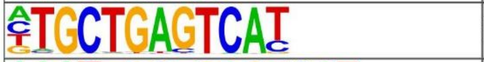 | Nrf2(bZIP)/Lymphoblast-Nrf2-ChIP-Seq(GSE37589)/Homer | 1e-10 | -2.365e+01 | 0.0000 | 16.0 | 3.44% | 173.2 | 0.36% |
| 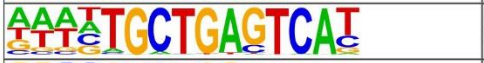 | Bach1(bZIP)/K562-Bach1-ChIP-Seq(GSE31477)/Homer | 1e-7 | -1.839e+01 | 0.0000 | 13.0 | 2.80% | 155.6 | 0.32% |
| 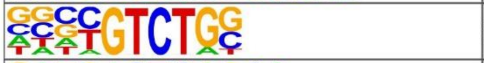 | Smad4(MAD)/ESC-SMAD4-ChIP-Seq(GSE29422)/Homer | 1e-6 | -1.578e+01 | 0.0000 | 124.0 | 26.67% | 8206.9 | 17.01% |
| 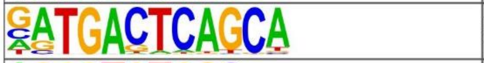 | NF-E2(bZIP)/K562-NFE2-ChIP-Seq(GSE31477)/Homer | 1e-5 | -1.365e+01 | 0.0000 | 12.0 | 2.58% | 201.7 | 0.42% |
| 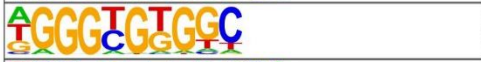 | KLF5(Zf)/LoVo-KLF5-ChIP-Seq(GSE49402)/Homer | 1e-5 | -1.333e+01 | 0.0000 | 128.0 | 27.53% | 8951.5 | 18.56% |
| 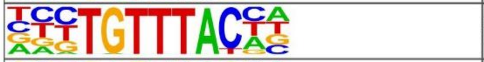 | FOXP1(Forkhead)/H9-FOXP1-ChIP-Seq(GSE31006)/Homer | 1e-5 | -1.234e+01 | 0.0001 | 32.0 | 6.88% | 1342.4 | 2.78% |
| 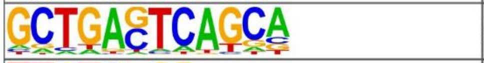 | MafK(bZIP)/C2C12-MafK-ChIP-Seq(GSE36030)/Homer | 1e-5 | -1.167e+01 | 0.0001 | 25.0 | 5.38% | 940.2 | 1.95% |
| 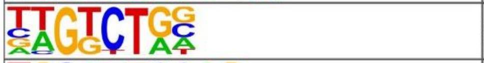 | Smad3(MAD)/NPC-Smad3-ChIP-Seq(GSE36673)/Homer | 1e-4 | -1.096e+01 | 0.0003 | 176.0 | 37.85% | 13886.7 | 28.79% |
| 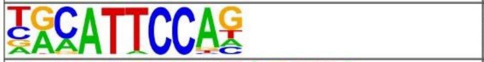 | TEAD3(TEA)/HepG2-TEAD3-ChIP-Seq(Encode)/Homer | 1e-4 | -1.067e+01 | 0.0003 | 72.0 | 15.48% | 4538.9 | 9.41% |
| 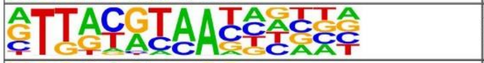 | NFIL3(bZIP)/HepG2-NFIL3-ChIP-Seq(Encode)/Homer | 1e-4 | -1.010e+01 | 0.0006 | 39.0 | 8.39% | 2009.8 | 4.17% |
| 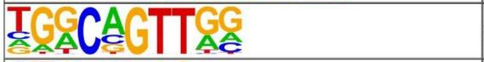 | AMYB(HTH)/Testes-AMYB-ChIP-Seq(GSE44588)/Homer | 1e-4 | -9.920e+00 | 0.0007 | 92.0 | 19.78% | 6359.0 | 13.18% |
| 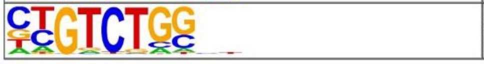 | Smad2(MAD)/ES-SMAD2-ChIP-Seq(GSE29422)/Homer | 1e-4 | -9.853e+00 | 0.0007 | 110.0 | 23.66% | 7968.3 | 16.52% |

### Motif enrichment in G12Vp53 ChIP peaks

Total Target Sequences = 2866, Total Background Sequences (genome) = 45361

| Motif | Name | P-value | log P-pvalue | q-value (Benjamini) | # Target Sequences with Motif | % of Targets Sequences with Motif | # Background Sequences with Motif | % of Background Sequences with Motif |
|---|---|---|---|---|---|---|---|---|
| 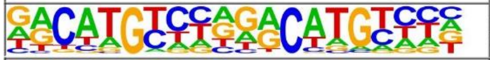 | p53(p53)/Saos-p53-ChIP-Seq(GSE15780)/Homer | 1e-2903 | -6.685e+03 | 0.0000 | 2316.0 | 80.81% | 440.0 | 0.97% |
| 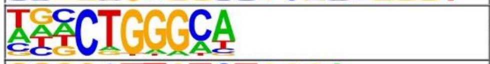 | ZNF416(Zf)/HEK293-ZNF416.GFP-ChIP-Seq(GSE58341)/Homer | 1e-186 | -4.290e+02 | 0.0000 | 1285.0 | 44.84% | 8973.8 | 19.81% |
| 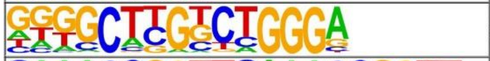 | Zfp809(Zf)/ES-Zfp809-ChIP-Seq(GSE70799)/Homer | 1e-97 | -2.244e+02 | 0.0000 | 303.0 | 10.57% | 993.5 | 2.19% |
| 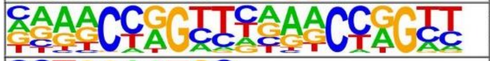 | Tcfcp2l1(CP2)/mES-Tcfcp2l1-ChIP-Seq(GSE11431)/Homer | 1e-85 | -1.980e+02 | 0.0000 | 208.0 | 7.26% | 498.3 | 1.10% |
| 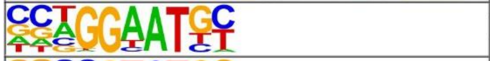 | TEAD4(TEA)/Tropoblast-Tead4-ChIP-Seq(GSE37350)/Homer | 1e-69 | -1.601e+02 | 0.0000 | 564.0 | 19.68% | 3892.9 | 8.59% |
| 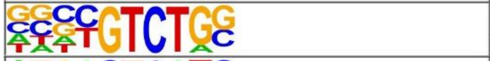 | Smad4(MAD)/ESC-SMAD4-ChIP-Seq(GSE29422)/Homer | 1e-62 | -1.432e+02 | 0.0000 | 817.0 | 28.51% | 7102.8 | 15.68% |
| 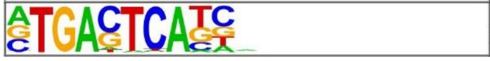 | AP-1(bZIP)/ThioMac-PU.1-ChIP-Seq(GSE21512)/Homer | 1e-47 | -1.104e+02 | 0.0000 | 405.0 | 14.13% | 2809.3 | 6.20% |

### Motif Enrichment in MCF10A-specific p53 ChIP peaks

Total Target Sequences = 565, Total Background Sequences (genome) = 46479

| Motif | Name | P-value | log P-pvalue | q-value (Benjamini) | # Target Sequences with Motif | % of Targets Sequences with Motif | # Background Sequences with Motif | % of Background Sequences with Motif |
|---|---|---|---|---|---|---|---|---|
| 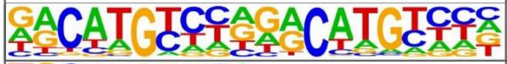 | p53(p53)/Saos-p53-ChIP-Seq(GSE15780)/Homer | 1e-871 | -2.006e+03 | 0.0000 | 523.0 | 92.57% | 523.2 | 1.11% |
| 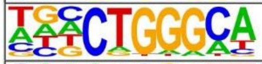 | ZNF416(Zf)/HEK293-ZNF416.GFP-ChIP-Seq(GSE58341)/Homer | 1e-34 | -7.939e+01 | 0.0000 | 222.0 | 39.29% | 8038.3 | 17.13% |
| 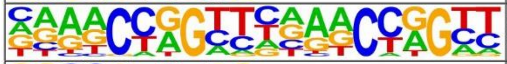 | Tcfcp2l1(CP2)/mES-Tcfcp2l1-ChIP-Seq(GSE11431)/Homer | 1e-27 | -6.268e+01 | 0.0000 | 47.0 | 8.32% | 446.9 | 0.95% |
| 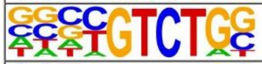 | Smad4(MAD)/ESC-SMAD4-ChIP-Seq(GSE29422)/Homer | 1e-16 | -3.792e+01 | 0.0000 | 165.0 | 29.20% | 7121.3 | 15.18% |
| 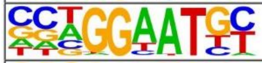 | TEAD4(TEA)/Tropoblast-Tead4-ChIP-Seq(GSE37350)/Homer | 1e-13 | -3.223e+01 | 0.0000 | 109.0 | 19.29% | 4115.8 | 8.77% |
| 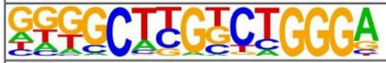 | Zfp809(Zf)/ES-Zfp809-ChIP-Seq(GSE70799)/Homer | 1e-11 | -2.580e+01 | 0.0000 | 35.0 | 6.19% | 700.6 | 1.49% |
| 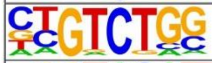 | Smad2(MAD)/ES-SMAD2-ChIP-Seq(GSE29422)/Homer | 1e-9 | -2.267e+01 | 0.0000 | 142.0 | 25.13% | 6954.1 | 14.82% |
| 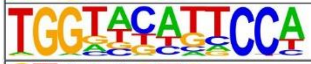 | PRDM10(Zf)/HEK293-PRDM10.eGFP-ChIP-Seq(Encode)/Homer | 1e-4 | -1.108e+01 | 0.0005 | 64.0 | 11.33% | 3046.5 | 6.49% |
| 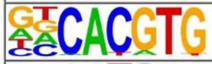 | BMAL1(bHLH)/Liver-Bmal1-ChIP-Seq(GSE39860)/Homer | 1e-4 | -1.102e+01 | 0.0005 | 162.0 | 28.67% | 9922.2 | 21.14% |
| 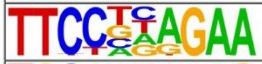 | STAT6(Stat)/Macrophage-Stat6-ChIP-Seq(GSE38377)/Homer | 1e-4 | -1.008e+01 | 0.0010 | 50.0 | 8.85% | 2264.5 | 4.83% |
| 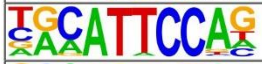 | TEAD3(TEA)/HepG2-TEAD3-ChIP-Seq(Encode)/Homer | 1e-4 | -9.612e+00 | 0.0016 | 98.0 | 17.35% | 5517.0 | 11.76% |
| 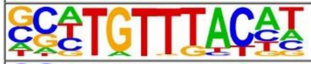 | FOXK2(Forkhead)/U2OS-FOXK2-ChIP-Seq(E-MTAB-2204)/Homer | 1e-3 | -7.545e+00 | 0.0116 | 58.0 | 10.27% | 3057.9 | 6.52% |
| 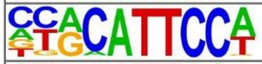 | TEAD1(TEAD)/HepG2-TEAD1-ChIP-Seq(Encode)/Homer | 1e-3 | -7.028e+00 | 0.0185 | 78.0 | 13.81% | 4511.4 | 9.61% |
| 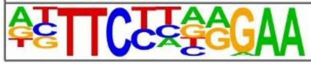 | STAT6(Stat)/CD4-Stat6-ChIP-Seq(GSE22104)/Homer | 1e-3 | -6.911e+00 | 0.0197 | 45.0 | 7.96% | 2275.0 | 4.85% |

**Supplementary Figure S4. Motif enrichment in cell type-specific p53 binding sites.**

Enrichment was calculated in G12V-specific, the whole G12V, and MCF10A-specific p53 binding loci (ChIP peaks) against the genomic sequence background.

**Supplementary Table S1. Differentially expressed genes.**

Differential gene expression between MCF10A and G12V cells based on two replicas of RNA-seq. First five columns indicate gene annotation- genomic location (chromosome, strand, transcription start and end site) and name. Next 4 columns indicate the CPM value in the two cell types in two replicas. The last three columns indicate log2 fold change, p Value and the direction of change: 1 and (-1) are up-regulated and down-regulated in G12V MCF10A cells respectively.

**Supplementary Table S2. Inverse primers used for 4C library amplification.**

| **Inverse PCR primers for 4C** | | |
|---|---|---|
| **Bait** | **Forward primer** | **Reverse primer** |
| FN1 | CCTGTGGCAATTACTGAAGGTG | GCAAGATAAACCTGTAGCTAAAGC |
| CDH1 | GCAGGGGCTAGAAACAAGC | ATTGTCACAACGATAAGGCC |
| FBN1 | TGCCACCTTTCAGAGACCAT | GGCATCCTGTTAAGCAACAAAG |
| TP53 | CAGCTGAGAGCAAACGCAAA | GGCTTTCCGTAATATCACACCC |
| MMP1 | CCATAAACAAGTCATTCTCAAAGC | ACTAAAGCCAAGATTTCTGTTACTG |

**Supplementary Table S3. Primers used for measuring gene expression by real time PCR.**

| **Gene** | **Primer 5' to 3'** | |
|---|---|---|
| **SDC1** | Forward | TCCCCACACAGAGGATGGAG |
| | Reverse | CTCCCCCGAGGTTTCAAAGG |
| **NDRG** | Forward | GTACTTCGTGCAGGGCATGG |
| | Reverse | GCGGGTGCCATCCAGAGAA |
| **COL17A1** | Forward | CTCCCCATCCCCAAGAAAGGC |
| | Reverse | GACCACACATGGGAGGGAAGG |
| **TP63** | Forward | GTCATTTGATTCGAGTAGAGGGG |
| | Reverse | CTGGGGTGGCTCATAAGGT |

| | | |
|---|---|---|
| **SERPINB5** | Forward | CGCAATGGATGCCCTGCAAC |
| | Reverse | AGGACATTGCCCAGTGGCTC |
| **EPS8L2** | Forward | CACACTGAGGTCTGCCCTTC |
| | Reverse | GTCCTTGGGGCTCATCTTGG |
| **EFNB1** | Forward | CCGCACACGCACCATGAAG |
| | Reverse | GCCTGTGTGGCCATCTTGAC |
| **TP53INP1** | Forward | GCCCAAGTAGTCCCAGAGTGG |
| | Reverse | CCACTGGGAAGGGCGAAAG |
| **FAM198B** | Forward | AGGATCCTGGGGCTCAACAG |
| | Reverse | GCGCATGGAAAGACACGCTG |
| **RGS12** | Forward | CCGGTCAACATCGACAGCC |
| | Reverse | TGGTACAGCGGGGACTTCAG |
